# Genome-wide variations analysis of special waxy sorghum cultivar Hongyingzi for brewing Moutai liquor

**DOI:** 10.1101/810580

**Authors:** Can Wang, Lingbo Zhou, Xu Gao, Yanqing Ding, Bin Cheng, Guobing Zhang, Ning Cao, Yan Xu, Mingbo Shao, Liyi Zhang

## Abstract

Hongyingzi is a special waxy sorghum (*Sorghum bicolor* L. Moench) cultivar for brewing Moutai liquor. For an overall understanding of the whole genome of Hongyingzi, we performed whole-genome resequencing technology with 56.10 X depth to reveal its comprehensive variations. Compared with the BTx623 reference genome, 2.48% of genome sequences were altered in the Hongyingzi genome. Among these alterations, there were 1885774 single nucleotide polymorphisms (SNPs), 309381 small fragments insertions and deletions (Indels), 31966 structural variations (SVs), and 217273 copy number variations (CNVs). These alterations conferred 29614 genes variations. It was also predicted that 35 genes variations were related to the multidrug and toxic efflux (MATE) transporter, chalcone synthase (CHS), ATPase isoform 10 (AHA10) transporter, dihydroflavonol-4-reductase (DFR), the laccase 15 (LAC15), flavonol 3′-hydroxylase (F3′H), flavanone 3-hydroxylase (F3H), *O*-methyltransferase (OMT), flavonoid 3′5′ hydroxylase (F3′5′H), UDP-glucose:sterol-glucosyltransferase (SGT), flavonol synthase (FLS), and chalcone isomerase (CHI) involved in the tannin synthesis. These results would provide theoretical supports for the molecular markers developments and gene function studies related to the liquor-making traits, and the genetic improvement of waxy sorghum based on the genome editing technology.

## Introduction

Sorghum [*Sorghum bicolor* (L.) Moench] is the fifth largest grain crop in the world after corn, wheat, rice, and barley, which is widely distributed in the arid and semi-arid regions of the tropics, and also one of the earliest cultivated cereal crops in China [1]. It has become a model crop for genome research of cereal crops because of its wide adaptability to environment, strong stress resistance, rich resources, and relatively small genome [2, 3]. According to different purposes, sorghum are generally divided into three types, namely sweet sorghum, feed sorghum, and grain sorghum. In grain sorghum, cultivars with amylose content between 0% and 5% are called waxy sorghum [4]. Waxy sorghum is one of the main raw materials for Moutai-flavor liquor and Luzhou-flavor liquor production due to its high amylopectin and tannin contents [5, 6]. In recent years, the undiversified main liquor-making waxy sorghum cultivar and its continuous degradation phenomenon has affected the supply of raw materials for liquor-making waxy sorghum and restricted the development of liquor enterprises [7]. Therefore, investigation of waxy sorghum genetic resources is a crucial measure for better straight evolution, genetic studies, and liquor-making waxy sorghum breeding strategies.

Genetic variation is a kind of variation that can be passed on to offspring due to the changes of genetic material in organisms and leads to the genetic diversity at different levels. There are many types of genetic variation in the genome, from microscopic chromosome inversion to single nucleotide mutation. With the development of genomics, the information of genetic variation that can be studied has become more comprehensive, such as single nucleotide polymorphism (SNP), small fragments insertion and deletion (Indel), structural variation (SV), and copy number variation (CNV) [8–10]. SNP is a kind of DNA sequence polymorphism caused by single base conversion or transversion, which is a new generation of molecular marker after restriction fragment length polymorphism (RFLP) and simple sequence repeats (SSR). It has been widely used in the construction of genetic linkage map, quantitative trait locus (QTL) mapping, genome-wide association study (GWAS), population genetic structure study, and genetic diversity analysis due to its characteristics of easy detection, large quantity, rich polymorphism, large flux, and wide distribution in genome [11–13]. Indel is a molecular biology term for an insertion or deletion of nucleotide fragments of different sizes at the same site in the genome sequence between the same or closely related species, which is widely distributed across the genome and occurs in a high density and large numbers in a genome. It has been applied to genetic analyses of animal and plant populations, molecular assisted crops and farmed animal breeding, human forensic genetics, and medical diagnostics because of its abundance, convenient typing platform, high accuracy, and good stability [14–16]. SV is operationally defined as genomic alterations that involve segments of DNA that are larger than 1 kb, and can be microscopic or submicroscopic, which mainly includes inversion, insertion, deletion, duplication, and other gene rearrangement. It can produce new genes, alter gene dosage and structure, and regulate gene expression elements, and have a significant impact on phenotypic variation and gene expression [17, 18]. CNV is a kind of genomic structural variation originated from gain or loss of DNA segments larger than 1 kb caused by genomic rearrangement, which has been reported to be associated with human complex diseases and widely used for prevention and clinical diagnoses of human diseases since it was first discovered in human populations. It is also widely found in the plant genomes, such as *Arabidopsis*, rice, corn, soybean, wheat, and cucumber, and its own gained or lost copies may result in the alteration of gene dosage and abundance of its transcript, and thus lead to the significant phenotypic variation of height, flowering time, and dormancy in plants [19, 20]. With the rapid development of molecular biology, whole-genome resequencing technology has been applied to genome-wide variations analysis in *Arabidopsis*, rice, maize, tomato, and other plants [8, 21–23]. The whole genome sequences of grain sorghum cultivar BTx623 has provided a template for genome-wide variations analysis in sorghum [24], and the first genome-wide variations analysis of sorghum was reported by [25].

Hongyingzi, a special waxy sorghum cultivar for brewing Moutai liquor containing 83.40% total starch, 80.29% amylopectin/total starch ratio, and 1.61% tannin. The genome-wide variation of Hongyingzi is not fully understood, yet it is necessary for liquor-making waxy sorghum functional genomic research and breeding. Here, we used whole-genome resequencing technology to study the whole genome variation of Hongyingzi, and discovered potential genome regions and metabolic pathways associated with liquor-making traits.

## Materials and methods

### Plant materials and whole-genome resequencing

Two sorghum cultivars were used in this study. Hongyingzi, approved by the Guizhou Crop Cultivar Approval Committee (Guiyang, Guizhou Province, China) in 2008, is a medium maturity waxy sorghum cultivar special used for brewing Moutai liquor and developed by Renhuai Fengyuan Organic Sorghum Breeding Center at Guizhou, China in 2008 [26]. BTx623 is an excellent grain sorghum cultivar used for whole-genome sequencing by the Joint Genome Institute and for constructing several mapping populations [25, 27].

Sees of Hongyingzi were sterilized by soaking in 0.1% mercury dichloride for 15 min, and then rinsed with distilled water for ten times. Next, seeds were placed in a germination box lined with three layers of filter paper and added 15 mL distilled water. The germination box was placed in the RXZ-1000B artificial climate box for cultivating 10 days as following parameters settings, day/night temperature is 28°C/25°C, light/dark time is 12 h/12 h, humidity is 85%, and light intensity is 340 μmol m^−2^ s^−1^. The 10-day-old healthy seedlings were harvested for DNA extraction using the CTAB (Hexadecyl trimethyl ammonium bromide) buffer method [28]. The DNA purity was determined by 0.8% agarose gel 100 V electrophoresis for 40 min and DNA concentration was determined by Qubit® 2.0 fluorescent meter (Invitrogen, Carlsbad, USA). Following quality assessment, the genomic DNA was randomly broken into 350 bp fragments by Covaris ultrasonic crushing apparatus and DNA fragments were end repaired, added ployA tail, added sequencing connector, purification, and PCR amplification to complete the establishment of the library. The constructed library was used to paired-end PE150 sequencing on Illumina HiSeq 4000 sequencing platform. The BTx623 reference genome sequences were downloaded from the https://phytozome.jgi.doe.gov/pz/portal.html#!info?alias=Org_SbicolorRio_er.

### Genomic variation detection and annotation

Bioinformatics analysis was carried out by Beijing Novogene technology co., LTD (Beijing, China). The original image data generated by the sequencing machine were converted into sequence data via base calling (Illumina pipeline CASAVA v1.8.2) and then subjected to quality control (QC) procedure to remove unusable reads according to following criteria: the reads contain the Illumina library construction adapters, the reads contain more than 10% unknown bases (N bases), and one end of the read contain more than 50% of low quality bases (sequencing quality value ≤ 5). After filtration, sequencing reads were aligned to the BTx623 reference genome using BWA [29] with default parameters. Subsequent processing, including duplicate removal was proformed using SAMtools [30] and PICARD (http://picard.sourceforge.net). The raw SNP/Indel sets were called by SAMtools with the parameters as ‘-q1 -C50 -m2 -F0.002 -d1000’, and then filtered this sets using the following criteria: the mapping quality > 20 and the depth of the variate position > 4. BreakDancer [31] and CNVnator [32] were used for SV and CNV detections respectively. ANNOVAR [33] was used for functional annotation of variants. The UCSC known genes were used for gene and region annotations.

### Gene variation analysis

Using the BTx623 gene set as the reference, genes with non-synonymous SNPs and Indels in coding regions identified in the Hongyingzi were selected as the candidate gene set. These genes were then aligned to the Gene Ontology (GO) and Kyoto Encyclopedia of Genes and Genomes (KEGG) database using Blast for clustering analysis [34, 35].

## Results

### Genome-wide identification of genetic variations in Hongyingzi

The whole genome of Hongyingzi was resequenced using Illumina Genome Analyser sequencing technology. The genome size of the BTx623 reference genome is 732152042 bp. Resequencing yielded 45.84 Gb of raw data, which comprised 45.79 Gb of high quality clean data (Table 1). There was a high sequencing quality (Q20 ≥ 97.55%, Q30 ≥ 93.10) and the GC content was 44.30%. The results showed that 97.52% of the Hongyingzi genome sequences (297504853 mapped reads) were identical to BTx623, average depth 56.10 X, with 95.94% of coverage at 1 X and 94.17% of coverage at least 4 X (Table 2). With these reads and the information from the BTx623 reference genome, large quantities of SNPs, Indels, SVs, and CNVs were identified (Fig. 1). Compared with the BTx623 reference genome, we finally found 1885774 SNPs, 309381 Indels, 31966 SVs, and 217273 CNVs in Hongyingzi.

**Table 1.** Summary of resequencing data of Hongyingzi.

| Raw base (Gb) | Clean base (Gb) | Effect rete (%) | Error rate (%) | Q20 (%) | Q30 (%) | GC content (%) |
| --- | --- | --- | --- | --- | --- | --- |
| 45.84 | 45.79 | 99.82 | 0.03 | 97.55 | 93.10 | 44.30 |

**Table 2.** Sequence alignment of Hongyingzi to BTx623.

| Mapped reads | Total reads | Mapping rate (%) | Average depth (X) | Coverage at least 1 X (%) | Coverage at least 4 X (%) |
| --- | --- | --- | --- | --- | --- |
| 297504853 | 305064750 | 97.52 | 56.10 | 95.94 | 94.17 |

**Fig. 1.**
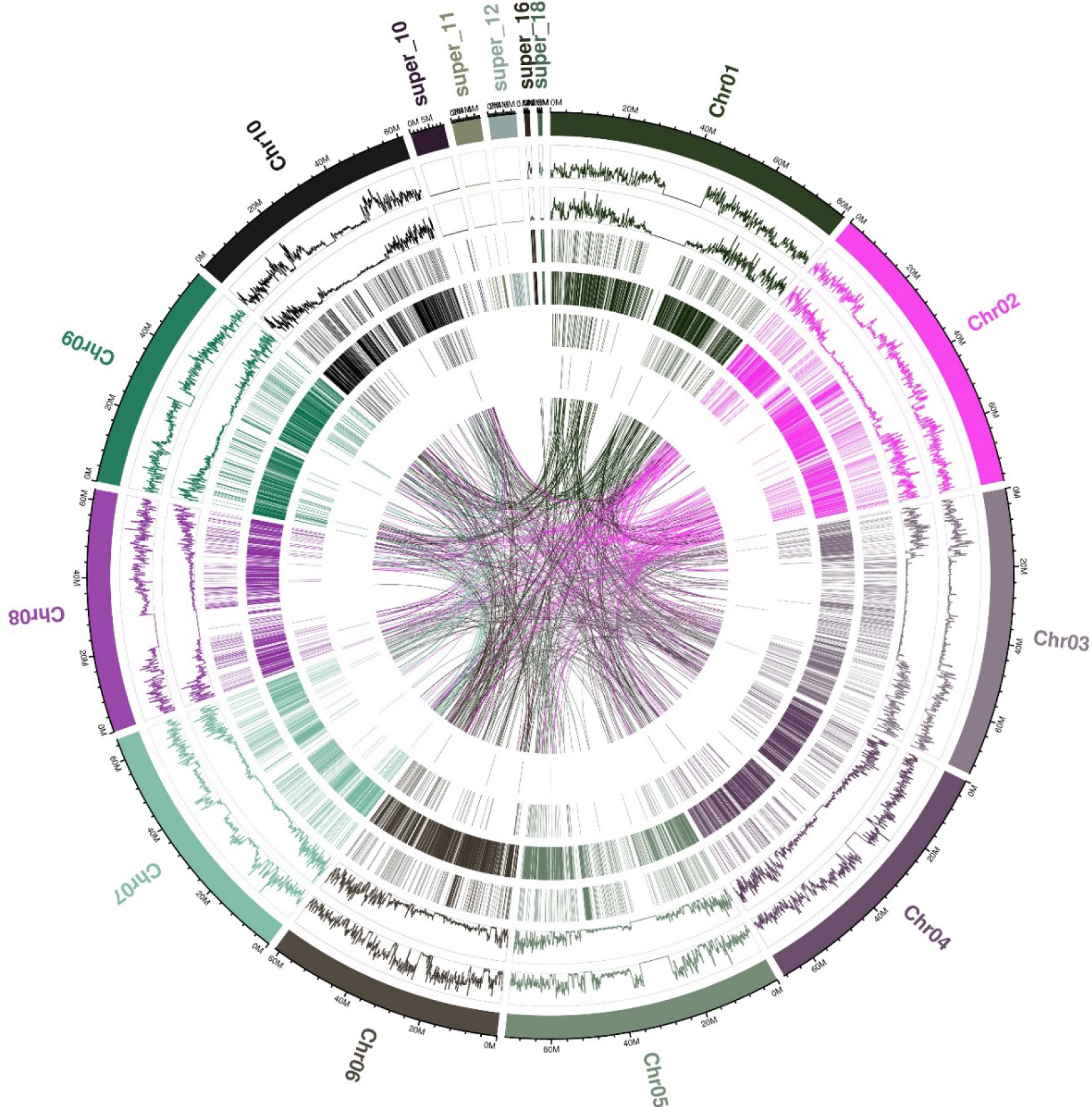
Genome-wide landscape of genetic variation in Hongyingzi. Cycles from outside to inside indicate chromosome, SNP, Indel, CNV duplication, CNV deletion, SV insertion, SV deletion, SV invertion, SV ITX, and SV CTX. **ITX**: Intrachromosomal translocation. **CTX**: Interchromosomal translocation.

### SNPs in the Hongyingzi genome

A total of 1885774 SNPs were identified in the Hongyingzi genome, including 1230508 transitions and 655266 transversions (Fig. 2A). Besides, there were 1401089 homozygous SNPs and 484685 heterozygous SNPs (Fig. 2B), and the het rate was 0.066%. As shown by annotations of SNPs detected in Hongyingzi (Table 3), there were 1515993 SNPs mutation in intergenic, 89326 SNPs in 1 kb of upstream, 75170 SNPs in 1 kb of downstream, and 6344 SNPs mutated in both 1 kb of upstream and downstream. We found that 76528 SNPs were mutated in exonic, including 38176 synonymous SNPs, 37774 non-synonymous SNPs, 453 SNPs related to gain of stop codons, and 125 SNPs related to loss of stop codons. We also found that there were 122211 SNPs mutation in intronic and 202 SNPs in splicing sites. Besides, the proportion of C:G>T:A type was observed to be the highest (Fig. 3).

**Fig. 2.**
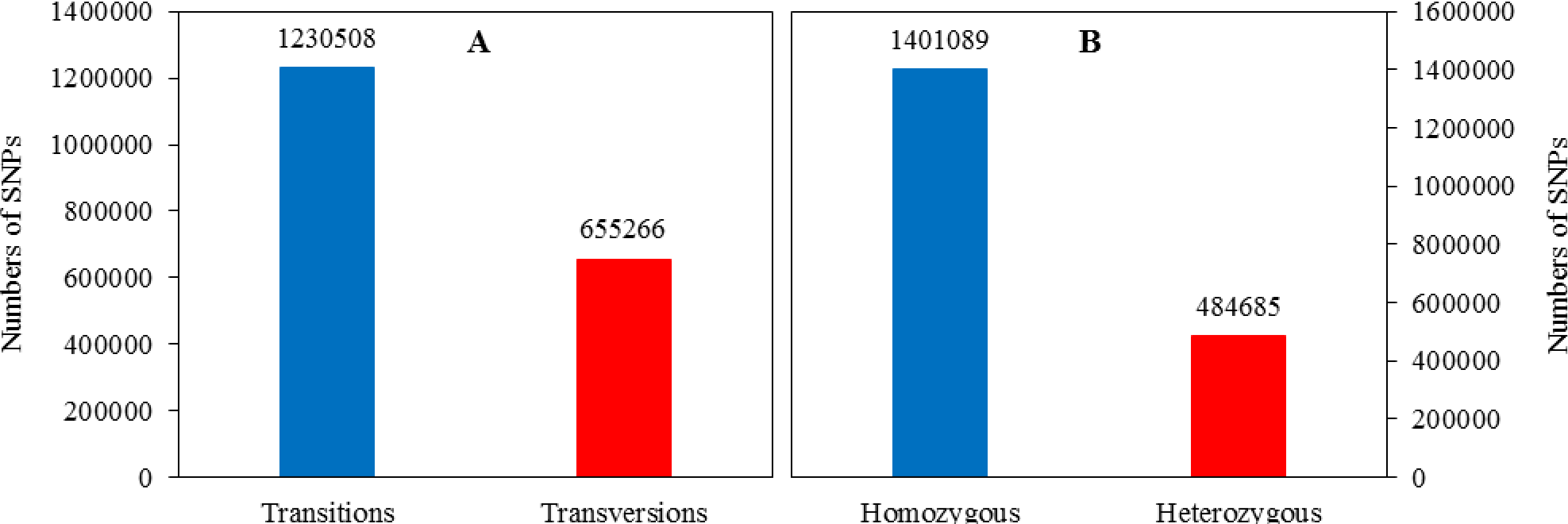
SNP distribution in the Hongyingzi genome. **A**: Numbers of transitions and transversions SNPs. **B**: Numbers of homozygous and heterozygous SNPs.

**Table 3.** Annotations of SNPs detected in Hongyingzi.

| Category | Numbers of SNPs | Region |
| --- | --- | --- |
| Intergenic | 1515993 |  |
| 1 kb of upstream | 89326 |  |
| 1 kb of downstream | 75170 |  |
| Both 1 kb of upstream and downstream | 6344 |  |
| Gain of stop codons | 453 | Coding regions |
| Loss of stop codons | 125 | Coding regions |
| Synonymous | 38176 | Coding regions |
| Non-synonymous | 37774 | Coding regions |
| Intronic | 122211 |  |
| Splicing sites | 202 |  |

**Fig. 3.**
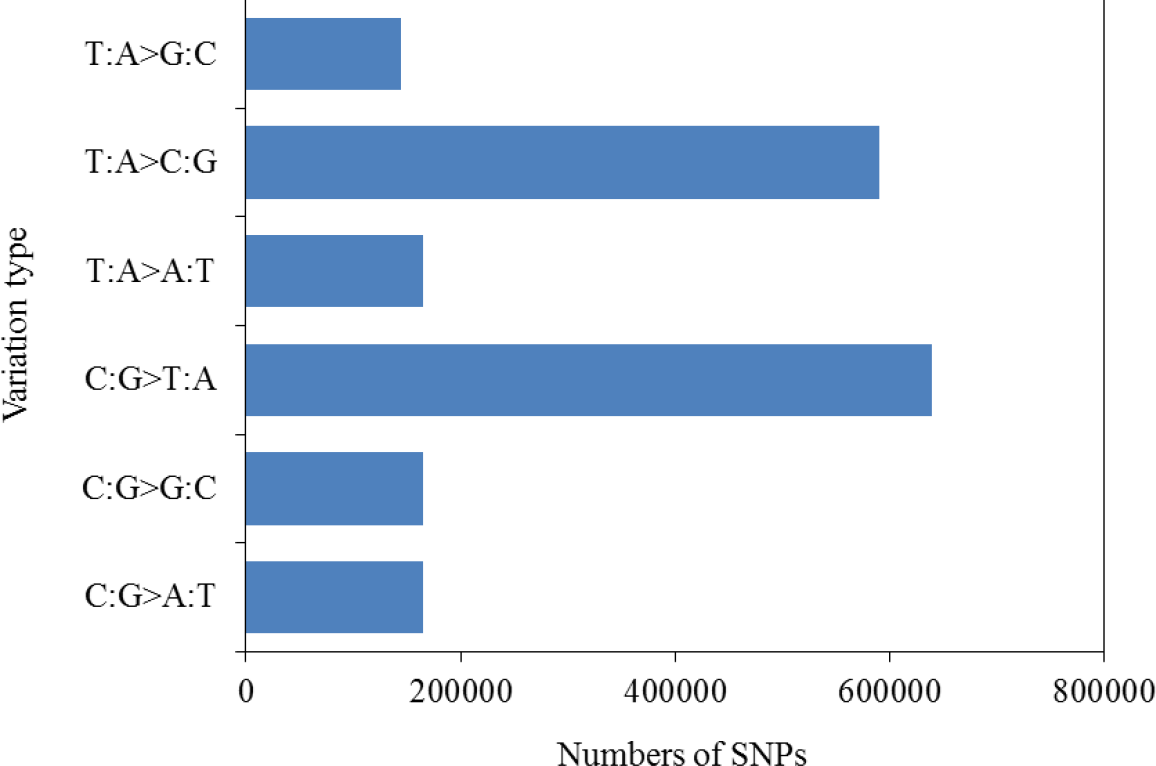
Distribution of SNP variation types

### Indels in the Hongyingzi genome

A total of 309381 Indels containing 149071 insertions and 160310 deletions, was uncovered in the Hongyingzi genome (Fig. 4A). These Indels also included 309361 homozygous and 20 heterozygous Indels (Fig. 4B), and the het rate was 0.0065%. Annotation analysis (Table 4) showed that there were 190165, 38198, 28361, and 2779 Indels mutated in intergenic, 1 kb of upstream, 1 kb of downstream, and both 1 kb of upstream and downstream, respectively. We found that 9375 Indels were mutated in exonic, in which 103 Indels were related to gain of stop codons, 22 Indels were related to loss of stop codons, 1354 insertions and 1476 deletions might lead to frameshift, and 3219 insertions and 3201 deletions might lead to non-frameshift. We also found 40223 Indels were mutated in intronic and 189 Indels did in splicing sites. Besides, the proportion of 1 bp (Fig. 5A) and 3 bp (Fig. 5B) Indels were observed to be the highest in whole genome and coding regions, respectively.

**Fig. 4.**
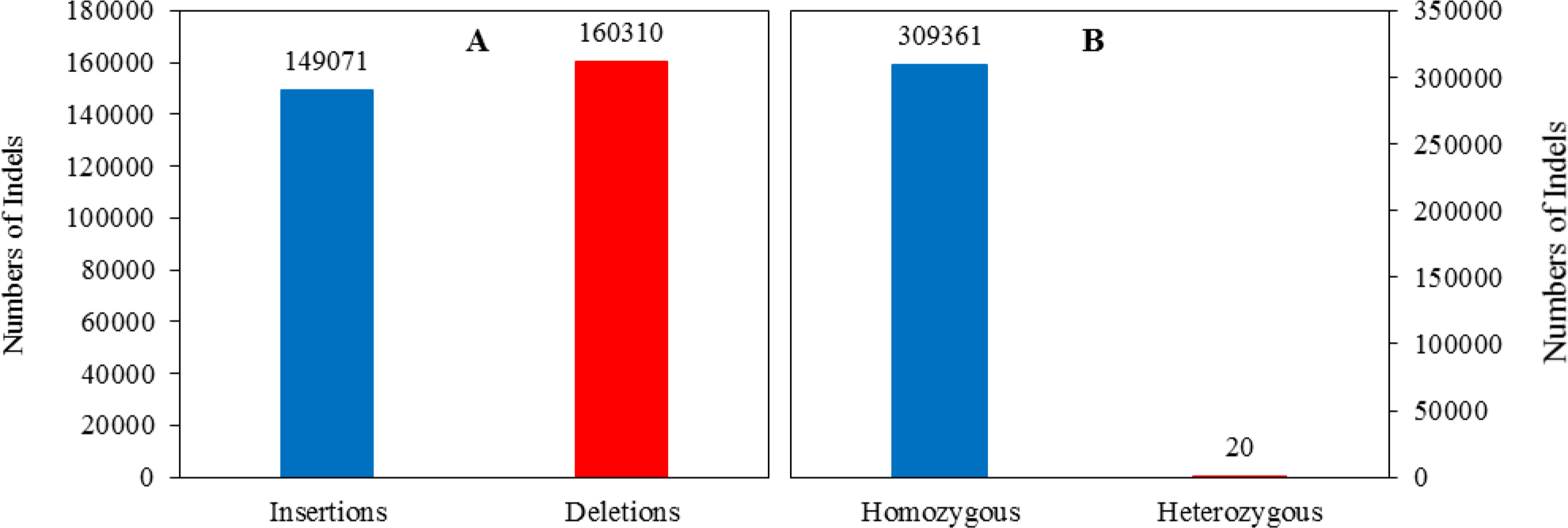
Indel distribution in the Hongyingzi genome. **A**: Numbers of insertions and deletions. **B**: Numbers of homozygous and heterozygous Indels.

**Table 4.**
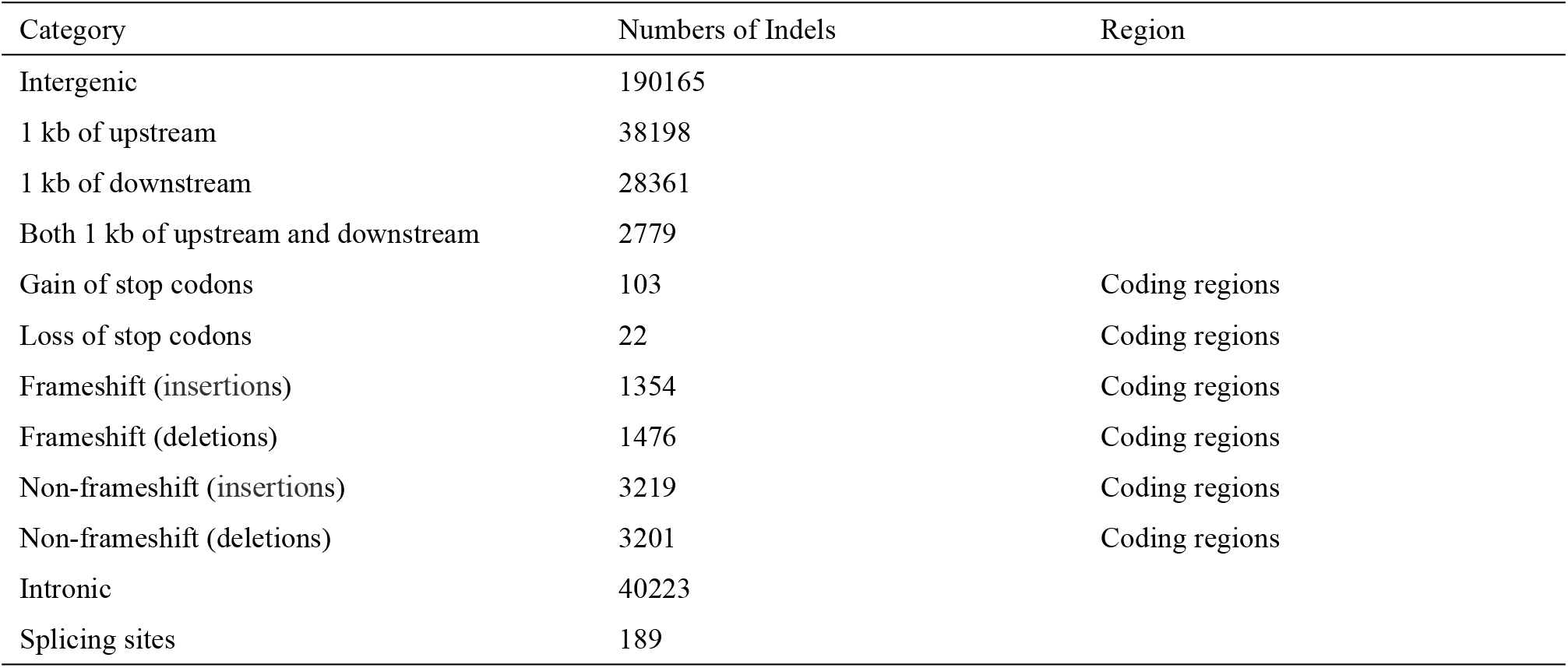
Annotations of Indels detected in Hongyingzi.

**Fig. 5.**
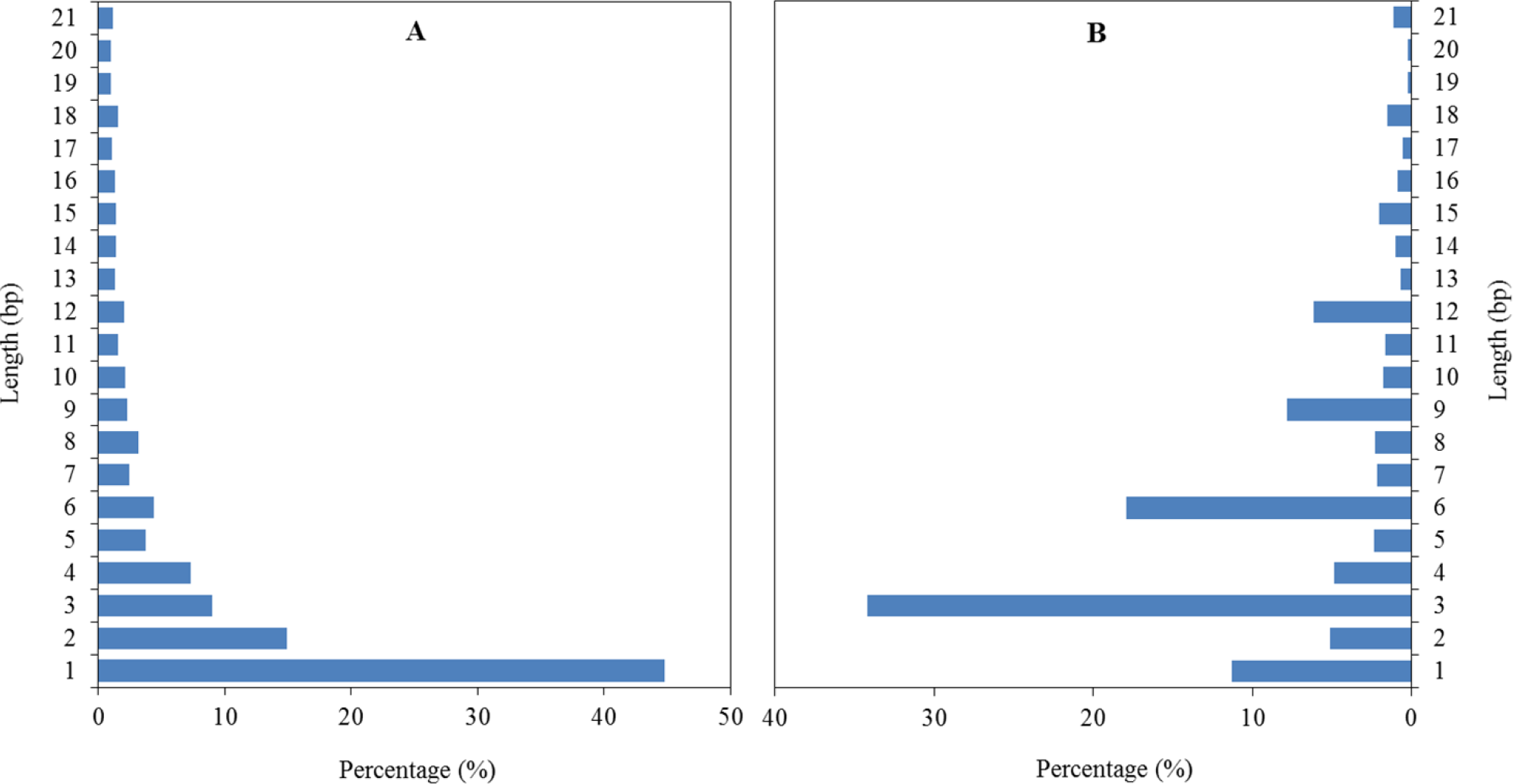
Length distribution of Indels in whole genome and coding regions. **A**: Length distribution of Indels in whole genome. **B**: Length distribution of Indels in coding regions.

### SVs in the Hongyingzi genome

A total of 31966 SVs were identified in the Hongyingzi genome, including 70 insertions, 15975 deletions, 1948 inversions, 4938 intrachromosomal translocations, and 9035 interchromosomal translocations (Fig. 6). As shown by annotations of SVs detected in Hongyingzi (Table 5), there were 9661 SVs mutation in intergenic, 1915 in 1 kb of upstream, 1460 in 1 kb of downstream, and 176 in both 1 kb of upstream and downstream. We also found that there were 3657 SVs mutation in exonic, 1119 in intronic, and 5 in splicing sites.

**Fig. 6.**
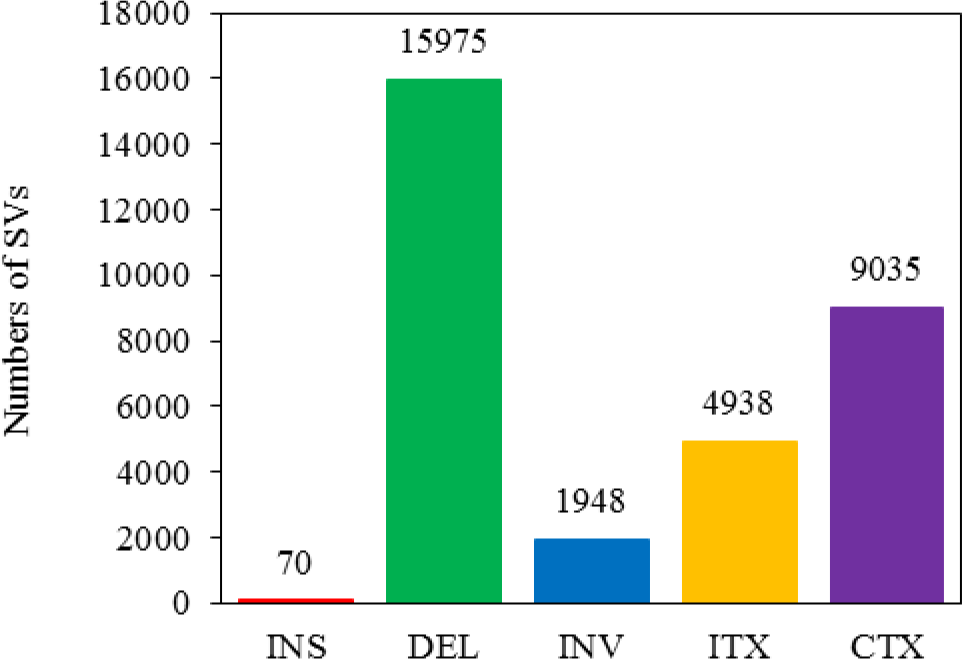
SV distribution in the Hongyingzi genome. **INS**: Insertions. **Del**: Deletions. **INV**: Inversions. **ITX**: Intrachromosomal translocations. **CTX**: Interchromosomal translocations.

**Table 5.** Annotations of SVs detected in Hongyingzi.

| Category | Numbers of SVs |
| --- | --- |
| Intergenic | 9661 |
| 1 kb of upstream | 1915 |
| 1 kb of downstream | 1460 |
| Both 1 kb of upstream and downstream | 176 |
| Exonic | 3657 |
| Intronic | 1119 |
| Splicing sites | 5 |

### CNVs in the Hongyingzi genome

A total of 217273 CNVs including 4966 duplications and 16307 deletions was uncovered in the Hongyingzi genome (Fig. 7). Annotation analysis (Table 6) showed that there were 17082, 985, 789, and 96 CNVs mutated in intergenic, 1 kb of upstream, 1 kb of downstream, and both 1 kb of upstream and downstream, respectively. We also found that there were 1822 CNVs and 496 CNVs mutated in exonic and intronic.

**Fig. 7.**
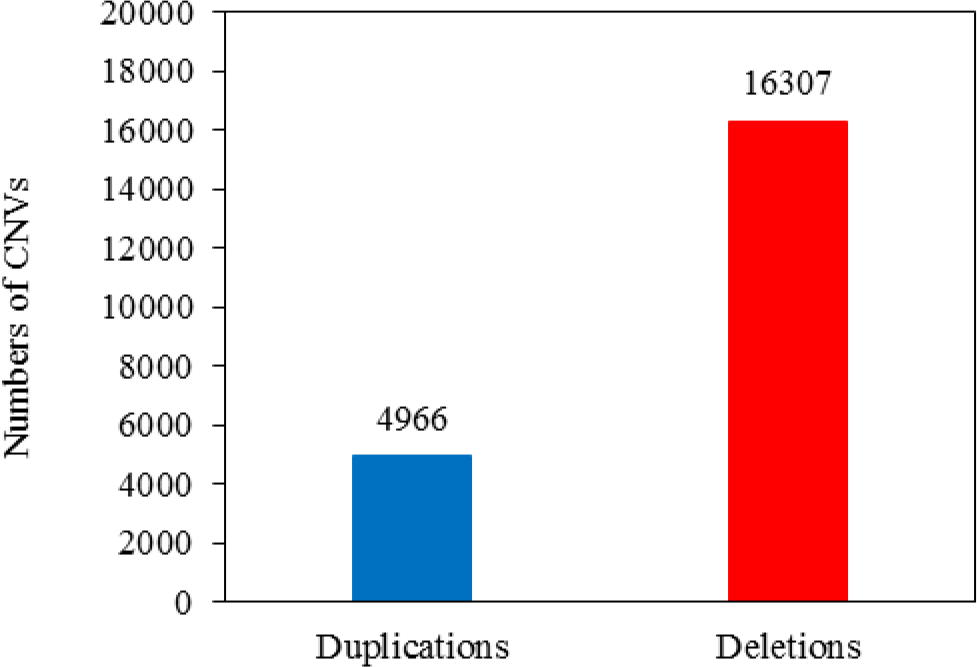
CNV distribution in the Hongyingzi genome.

**Table 6.** Annotations of CNVs detected in Hongyingzi.

| Category | Numbers of CNVs |
| --- | --- |
| Intergenic | 17082 |
| 1 kb of upstream | 985 |
| 1 kb of downstream | 789 |
| Both 1 kb of upstream and downstream | 96 |
| Exonic | 1822 |
| Intronic | 496 |

### Functional clustering of gene variations

Compared to the BTx623 reference genome, 29614 genes variations were identified in the Hongyingzi genome (Table 7). Of which, 14028, 25166 and 3948 was caused by SNPs, Indels, and 3948 SVs, respectively. GO annotation showed that SNPs and Indels were distributed among different gene ontologies (Fig. 8). In cellular component ontology, the cell and cell part contained the majority of gene variations with 19.06% SNPs and 23.01% Indels. Extracellular matrix contained a lower rate of variation. In molecular function ontology, binding and catalytic activity had a higher rate of variation. Binding included 40.77% and 37.56% of variation in SNPs and Indels, while catalytic activity did 34.01% 31.67% of variation in SNPs and Indels. In biological process ontology, metabolic process and cellular process had a high rate of variation. Metabolic process term included 39.00% and 36.07% of variation in SNPs and Indels, while cellular process did 39.05% and 35.99% of variation in SNPs and Indels. In KEGG annotation, 141 gene variations caused by SNPs (Fig. 9A) involved in the ubiquitin mediated proteolysis, while 1756 caused by Indels (Fig. 9B) involved in the metabolic pathways. These variations may affect the distinguishing traits between Hongyingzi and BTx623.

**Table 7.** Summary of gene variations in Hongyingzi.

| Variation types |  |  | Total |
| --- | --- | --- | --- |
| SNPs | Indels | SVs |  |
| 14028 | 25166 | 3948 | 29614 |

**Fig. 8.**
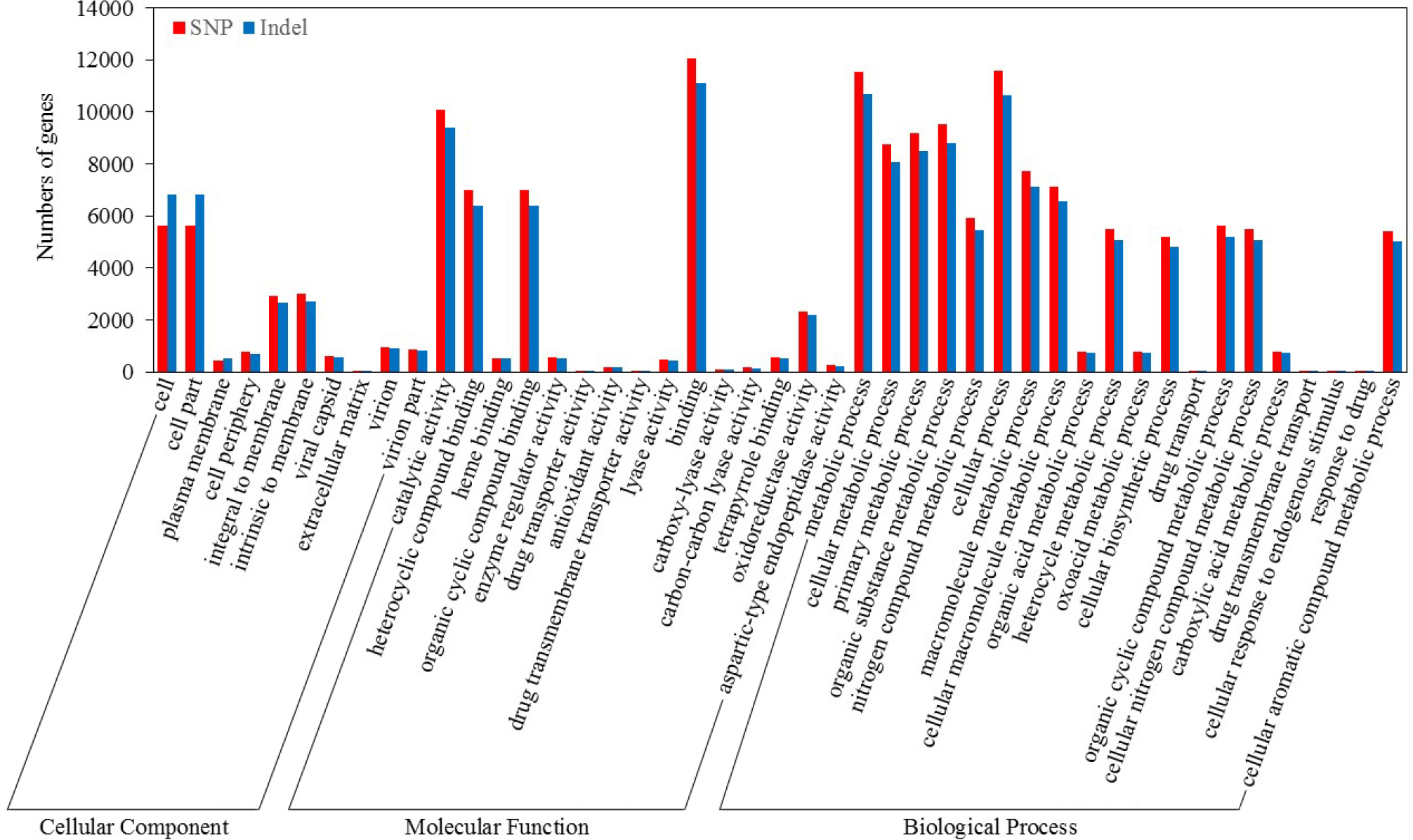
Classification of gene variations compared with GO database.

**Fig. 9.**
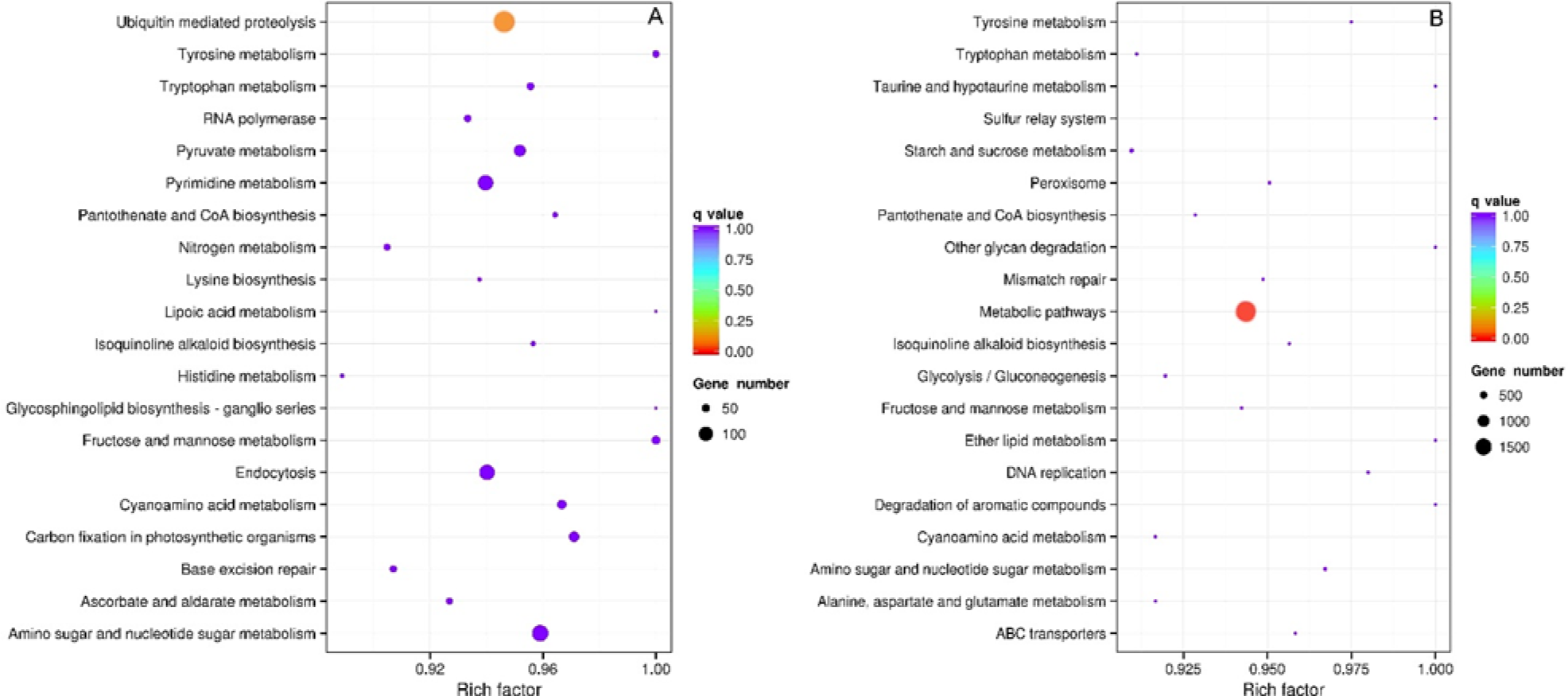
Classification of gene variations compared with KEGG database. **A**: Gene variations caused by SNPs. **B**: Gene variations caused by Indels.

### Genes variations involved in tannin synthesis

Compared to the BTx623 reference genome, we found that 35 genes variations were related to the tannin synthesis in the Hongyingzi genome (Table 8). Of which, 7 genes did in the multidrug and toxic efflux (MATE) transporter, 7 involved in the chalcone synthase (CHS), 4 did in the ATPase isoform 10 (AHA10) transporter, 4 did in the dihydroflavonol-4-reductase (DFR), 3 did in the laccase 15 (LAC15), 2 did in the flavonol 3′-hydroxylase (F3′H), 2 did in the flavanone 3-hydroxylase (F3H), 2 did in the *O*-methyltransferase (OMT), 1 did in the flavonoid 3′5′ hydroxylase (F3′5′H), 1 did in the UDP-glucose:sterol-glucosyltransferase (SGT), 1 did in the flavonol synthase (FLS), and 1 did in the chalcone isomerase (CHI).

**Table 8.**
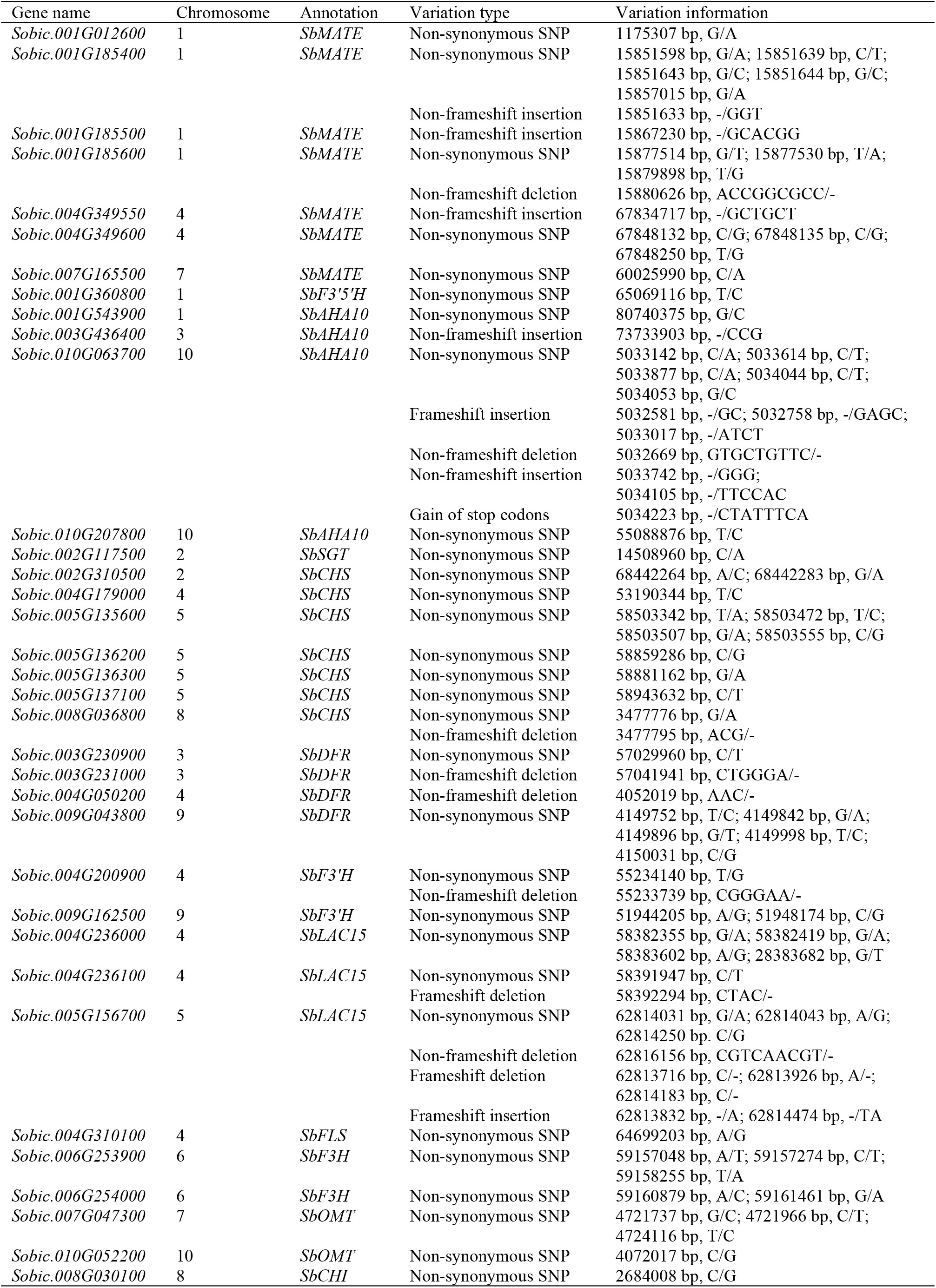
Summary of tannin synthesis related gene variations.

## Discussion

The rapid development of high-throughput sequencing technologies and bioinformatic tools makes it possible to understand the genetic variation and diversity of sorghum at the whole genome level, which plays an important role in enriching sorghum germplasm resources [24, 25, 36]. In this study, we used whole-genome resequencing technology to analyze the genetic variation in Hongyingzi, which is a special waxy sorghum cultivar for brewing Moutai liquor. The results showed that found that 2.48% of genome sequences were different between Hongyingzi and BTx623, and more than two million SNPs and Indels, along with large numbers of SVs and CNVs were identified. This is the first report on the genome-wide variations analysis in liquor-making waxy sorghum, which will be valuable for further genotype-phenotype studies and for molecular marker assisted breeding of liquor-making waxy sorghum.

In this study, the proportion of SNPs in intronic regions was 6.48%, which was higher than that in *Arabidopsis* [37]. Because the average intron size of sorghum is 444 bp, while the *Arabidopsis* is 168 bp [25]. A large number of SNPs was identified to alter in 202 splicing sites, 453 gain of stop codons, and 125 loss of stop codons. These alterations could lead to open reading frames extension, functional gene expression failure, or intron size increase [8, 21, 38]. Besides, the proportion of 3 bp Indels was observed to be the highest in coding regions. This might be due to the loss or increase of three bases results in the deletion or addition of a single amino acid without disrupting the overall reading frame [39], which could be a protection means to avoid the drastic changes of the genetic coding information, and then reduce damage to organisms due to natural variation. In addition, Indels with no multiples of 3 bp were rare in coding regions but relatively common in non-coding regions, because most of frameshift mutations is harmful to sorghum survival [25]. Compared to the BTx623 reference genome, a large number of SVs and CNVs was presented in the Hongyingzi genome, and the annotations of SVs and CNVs were similar to that of SNPs and Indels.

Compared to the BTx623 reference genome, there were 29614 genes variations in the Hongyingzi genome and Indels accounted for most of the genes variations. However, previous studies reported that SNPs accounted for most of the genes variations in *Arabidopsis* [40] and sorghum [25]. There are two possible reasons: 1) different materials used in different research, 2) limitations of early sequencing technology. Studies of SVs and CNVs in sorghum lag behind those in other plants. Recent studies in maize showed it potentially contributed to the heterosis during domestication and disease responses [41, 42]. Thus, we should focused on non-synonymous SNPs and Indels in coding regions for subsequent analysis of mutative genes. In our study, GO annotation showed that the mutative genes were equal distribution in different GO term. This indicates that SNPs and Indels may share similar survival and distribution patterns, although the origins and scales may different for affected genome segments.

Tannin, also known as condensed tannin or proanthocyanidins, is oligomers and polymers of flavan-3-ols [43, 44]. Sorghum has been the raw material for making famous liquor because of its grains containing tannin, and contributed special taste to Moutai-flavor liquor [45, 46]. Previous studies have mapped some gene loci associated with tannin content of sorghum. The *Tan1* gene (*Sb04g031730*) was cloned, which code a WD40 protein and control the tannin biosynthesis [43]. Two gene loci linked to tannin content were found [47]. One was named as *Sb01g001230*, coding glutathione-S-transferase, another was named as *Sb02g006390*, coding bHLH transcription factor and was isotopic with gene *B_2_* for color seed coat. Compared to the BTx623 reference genome, 35 genes variations were related to the tannin synthesis in the Hongyingzi genome. The genes involved in the MATE transporter, CHS, AHA10 transporter, DFR, LAC15, F3′H, F3H, OMT, F3′5′H, SGT, FLS, and CHI. Its variations would provide theoretical supports for the molecular markers developments and gene cloning, and the genetic improvement of waxy sorghum based on the genome editing technology.

## Conclusions

This is a first report of genome-wide variations analysis in liquor-making waxy sorghum. High-density SNP, Indel, SV, and CNV markers reported here will be a valuable resource for future gene-phenotype studies and the molecular breeding of liquor-making waxy sorghum. Genes variations involved in tannin synthesis reported here will provide theoretical basis for marker developing and gene cloning.

